# Spatial visualization of A-to-I Editing in cells using Endonuclease V Immunostaining Assay (EndoVIA)

**DOI:** 10.1101/2024.03.04.583344

**Authors:** Alexandria L. Quillin, Benoît Arnould, Steve D. Knutson, Jennifer M. Heemstra

## Abstract

Adenosine-to-Inosine (A-to-I) editing is one of the most widespread post-transcriptional RNA modifications and is catalyzed by adenosine deaminases acting on RNA (ADARs). Varying across tissue types, A-to-I editing is essential for numerous biological functions and dysregulation leads to autoimmune and neurological disorders, as well as cancer. Recent evidence has also revealed a link between RNA localization and A-to-I editing, yet understanding of the mechanisms underlying this relationship and its biological impact remains limited. Current methods rely primarily on *in vitro* characterization of extracted RNA that ultimately erases subcellular localization and cell-to-cell heterogeneity. To address these challenges, we have repurposed Endonuclease V (EndoV), a magnesium dependent ribonuclease that cleaves inosine bases in edited RNA, to selectively bind and detect A-to-I edited RNA in cells. The work herein introduces Endonuclease V Immunostaining Assay (EndoVIA), a workflow that provides spatial visualization of edited transcripts, enables rapid quantification of overall inosine abundance, and maps the landscape of A-to-I editing within the transcriptome at the nanoscopic level.

## Introduction

Adenosine-to-Inosine (A-to-I) editing is one of the most widespread RNA modifications in metazoans and is catalyzed by adenosine deaminases acting on RNA (ADARs).^1^ Among the three ADAR genes encoded in vertebrates (*ADAR1*, *ADAR2*, and *ADAR3*), ADAR1 is ubiquitously expressed and is responsible for the majority of editing in cells.^2,3^ ADAR1 is expressed in a p110 isoform that primarily resides in the nucleus, in addition to an IFN-inducible p150 isoform that localizes to the cytoplasm. Adenosine deamination results in a change of hydrogen bonding such that the resulting inosine base pairs with cytidine, effectively recoding the site to be recognized as guanosine by cellular machinery (Fig. 1a).^2^ Therefore, editing that occurs in protein-coding regions of messenger RNA (mRNA) can lead to multiple protein isoforms and altered function.^2^ While A-to-I editing is most abundant in non-coding, repetitive regions of mRNA and impacts transcript stability and localization,^2,4^ non-coding RNAs such as small-interfering RNAs (siRNAs) and microRNAs (miRNAs) are also edited, affecting global gene expression and cellular functionality.^5,6^ Collectively, A-to-I editing has proven essential for stem cell differentiation, embryogenesis, brain development, and cellular immunity.^2,7,8^

**Fig. 1.**
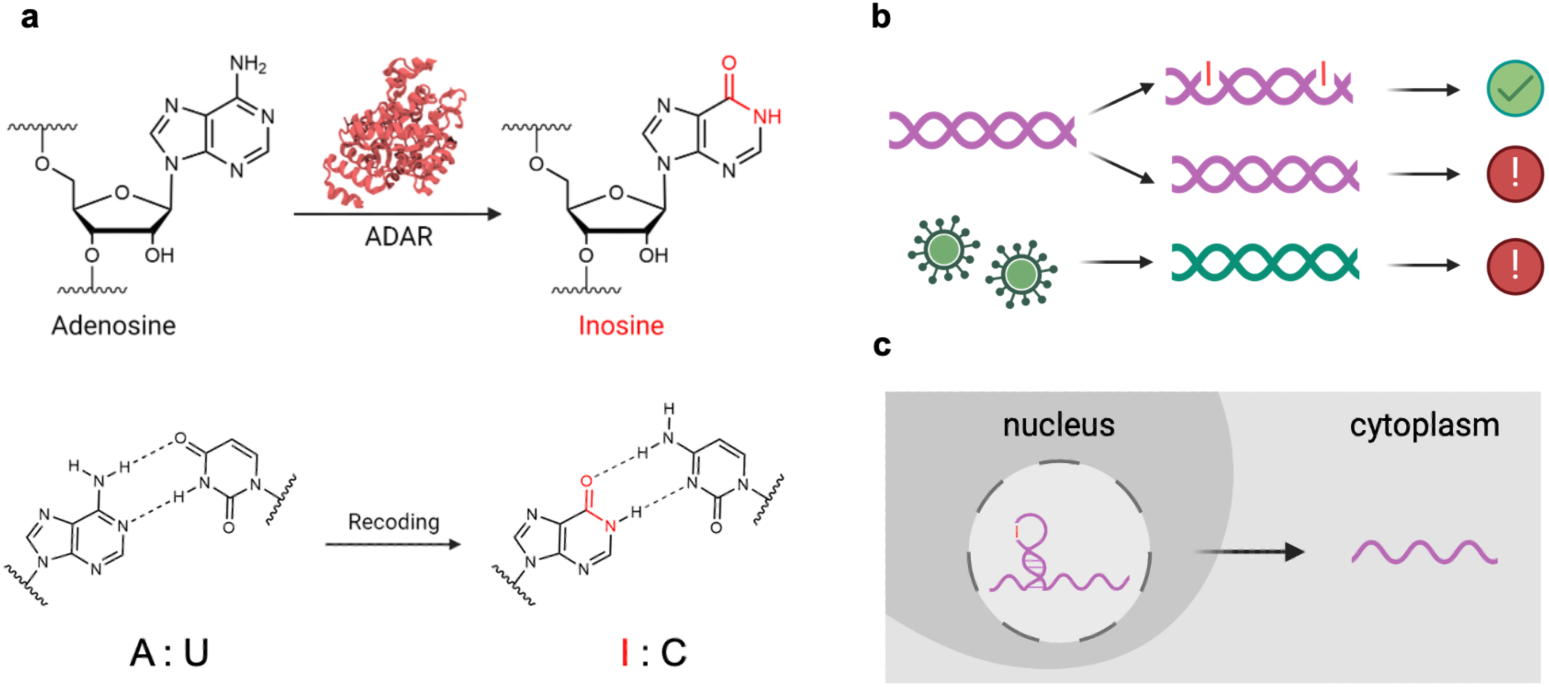
| A-to-I editing impacts essential biological functions. **a**, ADAR enzymes catalyze adenosine-to-inosine RNA editing, recoding edited sites to be read as guanosine to cellular machinery. **b,** Loss of A-to-I editing in endogenous RNAs triggers the innate immune response, synonymous to viral infections. **c,** Presence of extensively edited 3’ UTR in mRNA results in nuclear retention to paraspeckles.

ADAR1 is believed to play a critical role in regulating the activation of innate cellular immunity by targeting double-stranded RNA (dsRNA) made from long, inverted Alu repetitive elements that are located within introns and untranslated regions.^9^ These embedded Alu repeats make up 10% of the human genome and create ∼300 base pair long RNAs when transcribed. Due to their abundance, two neighboring Alu repeats can hybridize and form long endogenous dsRNAs that look very similar to foreign, viral RNA. ADAR prevents the activation of the cytosolic innate immune system by editing these endogenous dsRNAs and marking them as “self,” thus avoiding sensing by the MDA5-MAVS pathway that triggers interferon responses (Fig. 1b). These embedded Alu repeats make up 10% of the human genome, accounting for millions of edited sites and the majority of ADAR1 editing activity.^10–13^ Given the significant role A-to-I editing plays in maintaining cellular function, it’s unsurprising that dysregulation has been linked to multiple neurological disorders, autoimmune disorders, and cancers.^6,14,15^ A-to-I editing levels can display hyper- or hypo-editing depending on the specific cancer type.^16,17^ Notably, knockout of ADAR1 has been shown to be lethal in cell and animal models, and ADAR1 inhibition shows significant promise as a therapeutic strategy for treating cancer.^9,18–24^ Moreover, it has been observed that in melanoma, an A-to-I editing deficiency directly contributes to the melanoma metastatic phenotype.^25^ Collectively, these findings along with others highlight a clear link between A-to-I editing and the disease state, while speaking to the intricate relationship between editing and cancer that has yet to be fully elucidated. Leveraging this connection toward diagnostics and therapeutics will require greater understanding of the relationship between editing and disease progression, which has yet to be achieved with currently available methods.

The significance of A-to-I editing extends beyond sheer quantity; its influence on RNA localization is equally pivotal. Extensive A-to-I editing of inverted Alu repeats located within the 3’ UTR of mRNAs plays an essential role in nuclear retention^26–29^ and the presence or absence of an edited 3’ UTR has been demonstrated to drastically alter RNA subcellular localization (Fig. 1c). One example is the *mCAT2* gene that encodes both a protein-coding *mCAT2* mRNA, as well as its regulating partner, CTN-RNA. Prasanth et al. discovered that the CTN-RNA contains an edited 3’ UTR that mediates its nuclear localization. However, under stress conditions, CTN-RNA is post-transcriptionally cleaved at its 3’ UTR and exported to the cytoplasm for translation as well.^27^ Chen et al. observed similar results with the human gene, *Nicolin 1*. Nicolin 1 is expressed in multiple isoforms, but strikingly the only isoform that remains within the nucleus features an extensively edited, inverted Alu repeat in its 3’ UTR.^28^ From these and other studies, researchers propose that the inosine-specific RNA-binding protein (p54^nrb^) interacts with the edited dsRNA, mediating its retention in the nucleus.^26–29^ Most recently, the long non-coding RNA nuclear paraspeckle assembly transcript 1 (NEAT1) was determined to be essential for paraspeckle formation and RNA sequestration.^29^ Notably, 333 genes containing extensively edited inverted Alu elements in their 3’ UTR have been identified, suggesting A-to-I editing impacts the localization of far more RNAs than previously characterized.^28^ Beyond Alu repeats, the 3’ UTRs of RNAs also contain *cis*-regulatory elements, often referred to as “zipcodes,” that influence the localization of mRNA.^30,31^ These sequences serve as encoded cellular addresses that RNA-binding proteins (RBPs) interact with to facilitate trafficking.^32^ Considering that A-to-I editing primarily occurs within these non-coding regions, mRNA localization is subject to change based on the extent of editing. These examples underscore the profound connection between A-to-I editing and RNA localization. However, these findings also represent the extent of our current knowledge of the A-to-I editing landscape due to the lack of tools capable of mapping inosine within cells.

The most widespread technique used to characterize A-to-I editing is RNA sequencing (RNA-seq), in which RNA is extracted from cells, pooled, and subjected to high-throughput sequencing. Resulting sequences are compared to a reference genome and edited sites are identified from A-G transitions.^5^ While it remains the gold standard for identifying edited transcripts, unfortunately RNA sequencing along with other common methods requires RNA extraction that ultimately erases subcellular localization (Fig. 2a). Extraction also precludes the opportunity to compare cell-to-cell variation *in situ*, which is a hallmark of tumor heterogeneity. Mellis et al. demonstrated that the editing status of specific transcripts could be visualized in single mammalian cells using inosine fluorescence *in situ* hybridization (inoFISH). However, this method does not address the need for a generalizable approach to detect the number and localization of edited RNAs across the transcriptome.^33^ Antibodies could prove advantageous towards this goal, yet efforts to generate an anti-inosine antibody have fallen short, lacking selectivity and reproducibility.^34^ Taken together, a method for detecting and quantifying edited RNA *in situ* remains an unmet challenge.

**Fig. 2.**
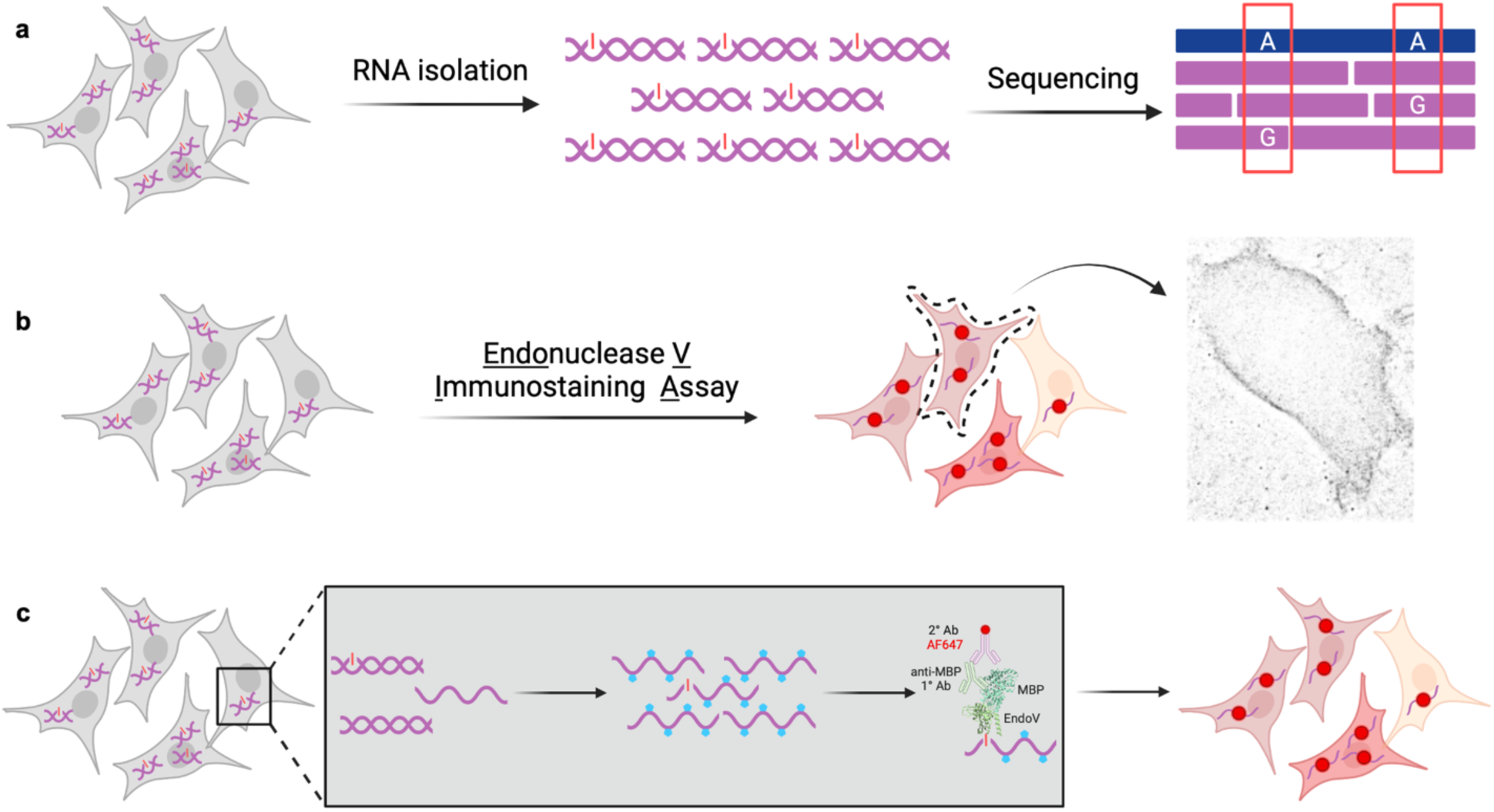
| Detection of inosine-containing RNA in cells using EndoVIA. **a**, RNA is isolated from cells and then sequenced to identify A-G transitions between RNA sequencing reads (purple) and a reference genome (blue), thus mapping A-to-I edited sites. **b,** EndoVIA detects cell-to-cell variation in A-to-I editing and localization of edited transcripts at the nanoscopic level. **c,** EndoVIA workflow; cells are fixed, treated with glyoxal (blue pentagons) to denature RNA secondary structure, prepped, and stained using EndoV and a series of antibodies to detect A-to-I edited sites *in situ*.

To address this need, we report herein Endonuclease V Immunostaining Assay (EndoVIA) which enables visualization of inosine-containing transcripts in cells using Endonuclease V (EndoV, Fig. 2b). EndoV is a magnesium-dependent ribonuclease that cleaves inosine-containing nucleic acid substrates; however, its activity can be controlled by replacing its natural magnesium cofactor with calcium such that EndoV binds inosine with high specificity without initiating cleavage.^35–38^ Our group has leveraged this binding event to ultimately repurpose EndoV as an “anti-inosine antibody” for enriching and quantifying edited RNA *in vitro*, which has in turn led to improved mapping of editing sites via RNA-seq as well as a microplate-based assay for quantifying global editing levels.^39,40^ Seeking to overcome the limitations of *in vitro* approaches, we recognized that the inosine binding capabilities of EndoV could be harnessed for detecting A-to-I edited transcripts *in situ* by recapitulating the principles of immunofluorescence (Fig. 2c). After optimization of our immunostaining protocol, we validated our EndoVIA workflow using cells with varying levels of A-to-I editing. We then demonstrated that our workflow could be used to detect both elevated levels of A-to-I editing in a hyper-editing cancerous cell line and reduced editing levels in a hypo-editing cancerous cell line. Finally, we used total internal reflection fluorescence (TIRF) illumination coupled with super-resolution microscopy to detect edited RNA at the single-molecule level. Excitingly, we observe distinct subcellular localization patterns for edited RNA in both healthy and diseased cell line models. Together, these results demonstrate that EndoVIA can provide spatial visualization of A-to-I editing, enable quantification of total cellular inosine content at the single-cell level, and be used to map subcellular localization of edited transcripts. This in turn provides a powerful tool for acquiring previously unattainable insights into the localization of A-to-I editing and its role in cellular processes and disease progression.

## Results

### Development of EndoVIA for *in situ* imaging of A-to-I Edited Transcripts

Seeking to advance our use of EndoV as an “anti-inosine antibody” and develop a method for detecting edited RNA *in situ*, we turned to immunofluorescence for inspiration. Immunofluorescence is a technique used to image cellular components by directly or indirectly labeling targets of interest with fluorophore-tagged antibodies.^41^ This approach has been used to label and image another abundant RNA modification, *N*^6^-methyladenosine (m^6^A), using an antibody specific for m^6^A-modified substrates. Through this approach, researchers were able to observe interactions with m^6^A-binding YTHDF proteins in stress granules.^42^ The immunofluorescence staining process involves cellular fixation, permeabilization, blocking, and staining with a primary antibody specific for the target, followed by a secondary antibody conjugated to a fluorophore.^41^ Using this basic protocol for immunostaining as a scaffold, we integrated essential steps necessary for imaging inosine-containing RNA with EndoV and optimized this workflow in a simple model system, human embryonic kidney 293T (HEK293T) cells.

Our first goal was to strategically fix cells in a way that maximizes staining specificity by preserving RNA integrity and eliminating cellular components that could lead to potential off-target binding. Fixation with reagents such as formaldehyde is used to preserve cellular morphology and prevent putrefaction.^41^ Although effective, formaldehyde crosslinks not only large RNAs such as ribosomal RNA (rRNA) and messenger RNA (mRNA), but also small RNAs including transfer RNA (tRNA) that are edited within the anticodon loop by adenosine deaminases acting on tRNA (ADATs).^43,44^ While essential for cell viability, tRNA editing is unrelated to ADAR-mediated A-to-I editing and tRNA is at least 100 times more abundant than mRNA in mammalian cells.^45^ Therefore, a significant concern was that fixation of tRNAs could mask our ability to visualize ADAR-edited targets. To circumvent this issue, we chose to fix cells using methanol (MeOH), another commonly used fixative reagent. Methanol preserves samples through dehydration and precipitation of proteins, retaining large RNAs and allowing small RNAs, like tRNA, to be washed away.^42,46^ To confirm that tRNA is indeed removed upon MeOH fixation, we performed tRNA FISH and observed that indeed the presence of tRNA dramatically decreased after fixation and washing (Supplementary Fig. 1). Organic solvents such as methanol have also been shown to retain nucleic acid content more effectively than aldehyde-based fixatives, which is essential to achieve the goal of our EndoVIA workflow.^46^

In addition to fixation method, we also explored additional workflow steps to achieve optimal EndoV binding of the target edited RNAs. Our group has shown that the commercially available, recombinant *Escherichia coli* EndoV binds edited single-stranded RNA (ssRNA) with a higher affinity than edited dsRNA.^39^ ADAR primarily targets RNA within double stranded regions, and thus an RNA denaturation step is required to maximize EndoV binding to ADAR1-edited sites.^1^ We were inspired by our previous use of glyoxal to denature RNA secondary structure while preserving EndoV binding capability *in vitro*, and hypothesized that we could recapitulate this approach in cells.^39^ Glyoxal is a reversible, chemical denaturant that forms bis-hemiaminal adducts with guanine, adenine, and cytosine nucleobases, thus disrupting Watson-Crick-Franklin base pairing.^39,40,47^ Importantly, glyoxal is unreactive towards inosine as inosine differs from guanine by the lack of an exocyclic amine that is critical for glyoxal adduct formation. Moreover, this small dialdehyde is also emerging as a popular fixative over classic reagents because of its efficiency and improved preservation of cellular morphology, providing further encouragement that this treatment would be compatible with our EndoV immunostaining protocol.^47^ Following methanol fixation and glyoxal treatment, cells were permeabilized with detergent and blocked with bovine serum albumin (BSA) to prevent nonspecific binding.

To achieve staining, cells were incubated sequentially with EndoV, primary antibody, fluorophore-labeled secondary antibody, all of which were diluted in a calcium-containing blocking buffer to support EndoV binding. The commercially available EndoV is fused to a maltose-binding protein (MBP) tag that is commonly used for purification purposes. Thus, we chose to use an anti-MBP primary antibody that binds MBP of the EndoV-MBP fusion protein. Accordingly, we utilized a secondary antibody that is specific for the primary anti-MBP antibody and is conjugated to Alexa Fluor 647 for visualization by fluorescence microscopy (Fig. 2c). Hoechst nuclear dye was also incorporated into the workflow to stain nuclear DNA as a point of reference.

### Optimization of EndoVIA

Our workflow as described is representative of typical immunofluorescence experiments, with some modifications to support use of EndoV as an “anti-inosine antibody.” To confirm that our modified workflow was suitable for immunofluorescence and that the modifications made to enable use of EndoV would not alter cellular morphology or interfere with imaging, we first sought to detect and image well-known and characterized cellular components using commercially available antibodies. We chose to stain β-actin, a highly abundant structure in the cytoskeleton, and nucleoporin 153 (Nup153), a less abundant protein located in nuclear pore complexes.^48,49,50,51^ Importantly, staining these controls would ensure that our immunostaining workflow has the dynamic range needed to capture the signal associated with edited RNA. β-actin and Nup153 were immunostained in HEK293T cells using the previously outlined workflow. As negative controls, cells were also stained by omitting the primary or secondary antibodies to assess nonspecific binding and autofluorescence. Images for both proteins were representative of previous reports, exhibiting sufficient signal and minimal background (Supplementary Fig. 2a and 2b).

Given that EndoV is fused to MBP, we considered the possibility that MBP could potentially engage in non-specific binding and give rise to unwanted background signal. To assess nonspecific binding of the fusion protein, we stained HEK293T cells initially with a 1:1000 dilution EndoV-MBP fusion or *Escherichia coli* MBP using our previously outlined staining workflow and imaged using widefield microscopy (Supplementary Fig. 3). As a negative control, a set of cells were not treated with EndoV fusion or MBP but were still incubated with primary and secondary antibody. Encouragingly, fluorescence was only observed for cells treated with EndoV-MBP, and cells treated with MBP only were comparable to negative controls. These results suggest the observed fluorescence is derived from the binding of EndoV to edited RNAs.

Detecting the maximum number of edited sites requires the complete denaturation of RNA secondary structure in order to grant EndoV access to these regions. To find the optimal glyoxal concentration for achieving this outcome, we stained *GAPDH* mRNA using FISH in HEK293T cells at varying concentrations of glyoxal (Supplementary Fig. 4). As RNA is increasingly glyoxylated, FISH probes are no longer able to hybridize to their targets, resulting in a decrease in fluorescence. We also stained β-actin in parallel under each glyoxal concentration to monitor cellular morphology (Supplementary Fig. 5). Taken together, we determined that 12% glyoxal was sufficient to denature RNA secondary structure without impacting cellular morphology.

Immunofluorescence protocols often require antibody optimization to minimize background and achieve maximum signal-to-noise ratio and sensitivity. Because we are using EndoV in a manner analogous to an antibody for inosine, optimizing concentration was thus imperative for developing a robust fluorescence-based assay that accurately detects edited substrates. To meet this need, we systematically screened EndoV concentrations to identify conditions where EndoV would stain the maximum number of targets while avoiding nonspecific binding of MBP. To satisfy these requirements, we leveraged the recombinant MBP used in the previous experiment to assess nonspecific binding once again. HEK293T cells were cultured and stained with increasing amounts of EndoV or MBP ranging from 1:4000 to 1:25, along with the appropriate controls (Supplementary Fig. 6). After quantifying the fluorescence, we determined that 1:50 EndoV was sufficient to achieve saturation while mitigating nonspecific binding of MBP. For further rigor, we also hypothesized that removing calcium, which is essential for EndoV binding, would greatly reduce the fluorescence of inosine-containing RNA. To test this, we treated cells with EDTA following EndoVIA and found that fluorescence significantly decreased, further supporting that the signal observed could be attributed to EndoV binding to edited RNA (Supplementary Fig. 7).

### Validation of EndoVIA

Having optimized our immunofluorescence protocol, we next sought to validate the ability of EndoVIA to detect biologically relevant differences in inosine by staining cells having varying levels of A-to-I editing. We predicted that if our workflow was targeting edited RNA as expected, then staining cells with little to no editing would lead to less fluorescence signal than staining of wildtype cells. To test our hypothesis, we chose to stain a HEK293T cell line with the ADAR1 gene knocked out via CRISPR-Cas9 genome editing.^9^ We first immunostained ADAR1 in both the WT and KO cell lines and confirmed that indeed ADAR1 was absent from the KO cell line (Supplementary Fig. 8). Using our optimized workflow, we then immunostained inosine-containing RNA in both wildtype and ADAR1 KO HEK293T cells and quantified the fluorescence (Fig 3a and 3b). Image analysis revealed that WT HEK293T cells show ∼2-fold greater fluorescence than ADAR1 KO cells, strongly suggesting that edited RNA is responsible for the staining.

**Fig. 3.**
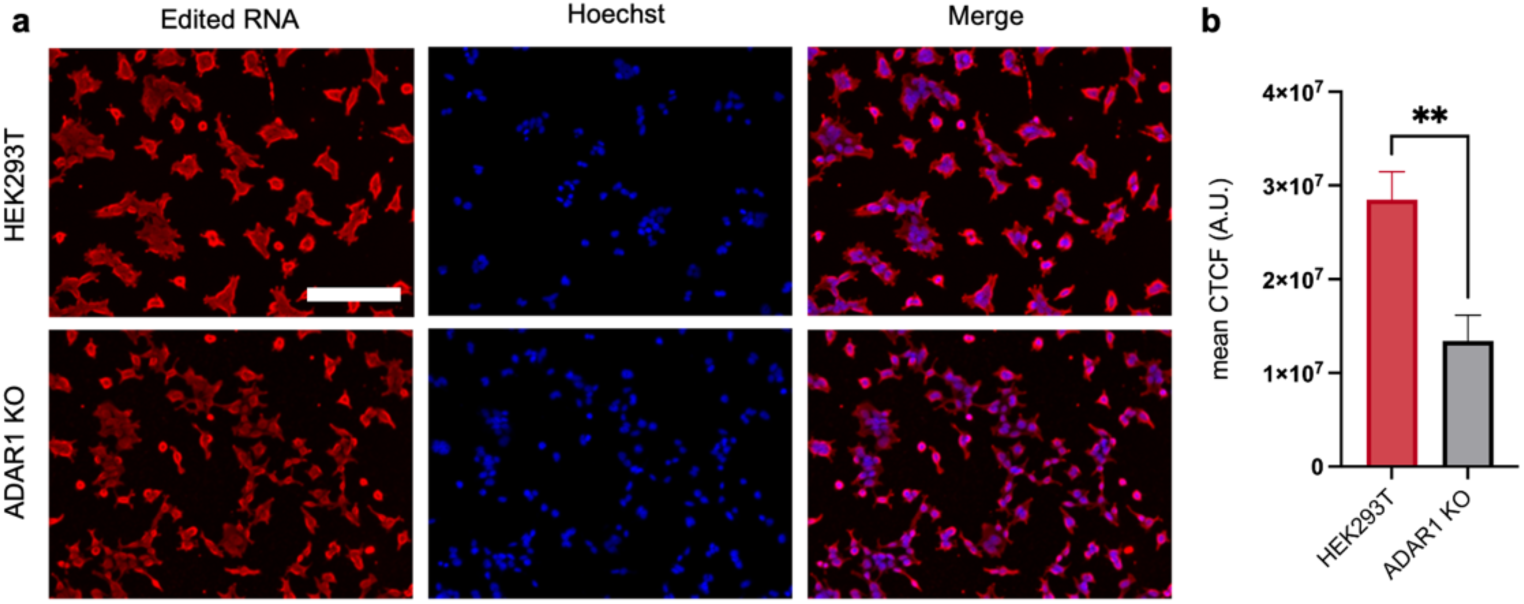
| Staining edited RNA in cells with varying levels of A-to-I editing. **a**, Fixed WT and ADAR1 KO HEK293T cells were immunostained using the optimized EndoVIA workflow for edited RNA (red) and cell nuclei (blue). **b,** Quantification of mean corrected total cellular fluorescence (CTCF) in **a**. Data in **a** and **b** are representative of three independent experiments; *n=3* wells from a 96-well plate. Scale bar, 200 μm. Data are shown as mean ± s.d. in arbitrary units (A.U.). Statistical significance was determined by unpaired *t*-test; \*\**P* < 0.01.

While we were encouraged by this significant difference, we were not entirely surprised by the remaining fluorescence from the ADAR1 KO cells. Other enzymes that are large contributors of cellular inosine, such as ADAR2 and ADAT, are still very much prevalent and may contribute to editing. We turned to RNA-seq and the Alu editing index (AEI) to compare the WT HEK293T and ADAR1 KO cells. The AEI was developed by Roth and coworkers and is a power computational tool that maps editing in Alu repeats across large datasets generated from RNA sequencing and has become widely accepted as the gold standard for quantifying ADAR editing activity.^12^ Briefly, the AEI value is generated from RNA-seq data by identifying sites that are called as guanine in RNA but adenosine in DNA and then calculating the ratio of these edited sites to the total coverage of adenosines. We determined that the AEI values for the WT HEK293T and ADAR1 KO cells were 1.4 and 0.2, respectively (Supplementary Fig. 9). These results suggest that there is some remaining edited tRNA or other unwanted species giving rise to background signal. Although the observed background signal was not ideal, we recognized that this could be especially noticeable in these very low-editing cells whereas it might not be as problematic in other cell lines and thus we carried forward. Encouragingly, our results described below indicate that this background signal appears to be nominal in the context of cells having more typical editing levels.

We were curious to explore whether we could detect an increase in A-to-I editing as well. Immortalized cell lines have low editing activity in comparison to human primary cells. Specifically, HEK293T cells have low editing levels, making them an ideal cell line to stimulate ADAR1 upregulation.^52^ To explore this hypothesis, we transfected HEK293T cells with increasing amounts of a GFP-tagged ADAR1 p110 overexpression vector (pCMV) plasmid and stained accordingly (Fig. 4a). We quantified the fluorescence of both ADAR-GFP expression and edited RNA and found that ADAR-GFP expression maximized at 100 ng of plasmid and decreased at higher concentrations. We also observed a direct relationship between the fluorescence of ADAR-GFP expression and edited RNA (Fig. 4b). To confirm that changes in fluorescence were due to ADAR transfection, we also transfected HEK293T cells with increasing amounts of coilin-GFP, another nuclear protein. We found that there was no change in edited RNA fluorescence (Supplementary Fig. 10). Previous literature has demonstrated that increasing ADAR1 expression does not necessarily lead to more A-to-I editing due to the presence of additional factors that regulate editing.^53,54^ Thus, we were encouraged to see a rough correlation between A-to-I editing fluorescence and ADAR1 levels but not surprised that the difference was far less clear (and not outside of error) compared to the ADAR1 KO cell line.

**Fig. 4.**
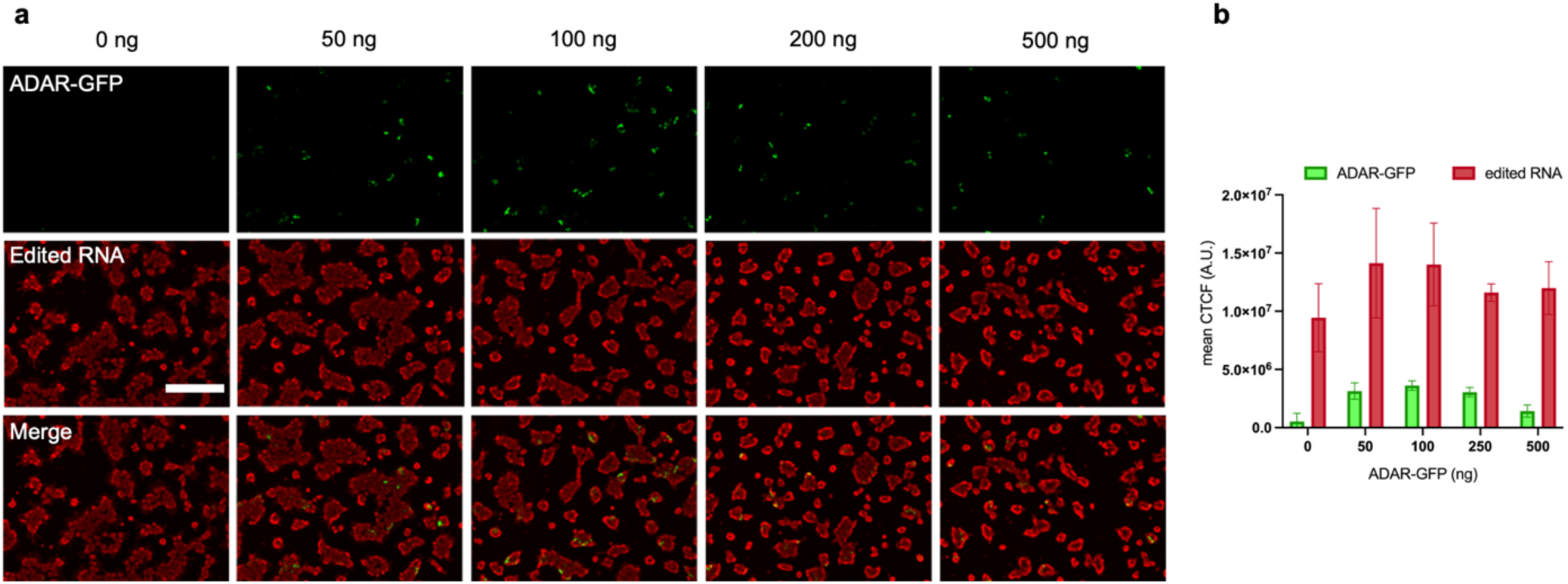
| Detecting an increase in A-to-I editing in HEK293T cells. **a**, HEK293T cells were transfected with increasing amounts of ADAR-GFP plasmid (0-500 ng, green), fixed, and stained for edited RNA (red) using EndoVIA. **b,** Quantification of mean corrected total cellular fluorescence (CTCF) of ADAR-GFP and edited RNA in **a**. Data in **a** and **b** are representative of three independent experiments; *n=3* wells from a 96-well plate. Scale bar, 200 μm. Data are shown as mean ± s.d. in arbitrary units (A.U.).

### Detecting A-to-I Editing in Cancer

Dysregulated A-to-I editing has been linked to various cancer types, and while many cancer types present a hyper-editing signature, some also display hypo-editing.^55^ Most notably, hyper-editing is evident in thyroid, head, neck, lung, and breast cancers whereas hypo-editing is observed in metastatic melanoma, invasive breast cancer, and renal cancer.^24,56–58^ As a result, A-to-I editing is rapidly emerging as not only a biomarker for diagnosing and studying cancers, but also as a potential therapeutic target. We recognized that EndoVIA could prove to be particularly powerful in this context, as it could enable rapid quantification of editing levels across multiple samples and with the ability to observe cell-to-cell heterogeneity. Additionally, we envisioned that the ability to discriminate between inosine levels in healthy vs cancerous cells could serve as a foundation for the development of phenotypic screening assays for ADAR inhibition.

Fortuitously, Schaffer et al. recognized the importance of having proper cellular models that reflect the A-to-I editing levels of a given tissue condition or disease signature and using the AEI computational tool, they determined the AEI values for more than 1,000 human cell line types.^59^ This provided a wealth of validated cell lines for us to choose from in exploring the ability of EndoVIA to detect cancer-related changes in inosine levels. Drawing inspiration from the Cell Line A-to-I Editing Catalogue, we first chose to test our workflow against ZR-75-1 and MCF10A cell lines. ZR-75-1 cells are a malignant breast cell line that displays hyper-editing signatures with an AEI value of 2.1 whereas the MCF10A breast cell line as a non-malignant control that has an AEI value of 1.4 (Fig. 5a). Both cell lines were subjected to our EndoVIA workflow, and we were very excited to see that the ZR-75-1 cell line did indeed show higher fluorescence than the MCF10A cells (Fig. 5b). Even more encouraging, the ratio between the immunofluorescence signal for the two cell lines was comparable to that of the reported AEI values. This demonstrates that EndoVIA is able to detect a cancer-related increase in RNA editing without the need for sequencing, which dramatically increases potential throughput.

**Fig. 5.**
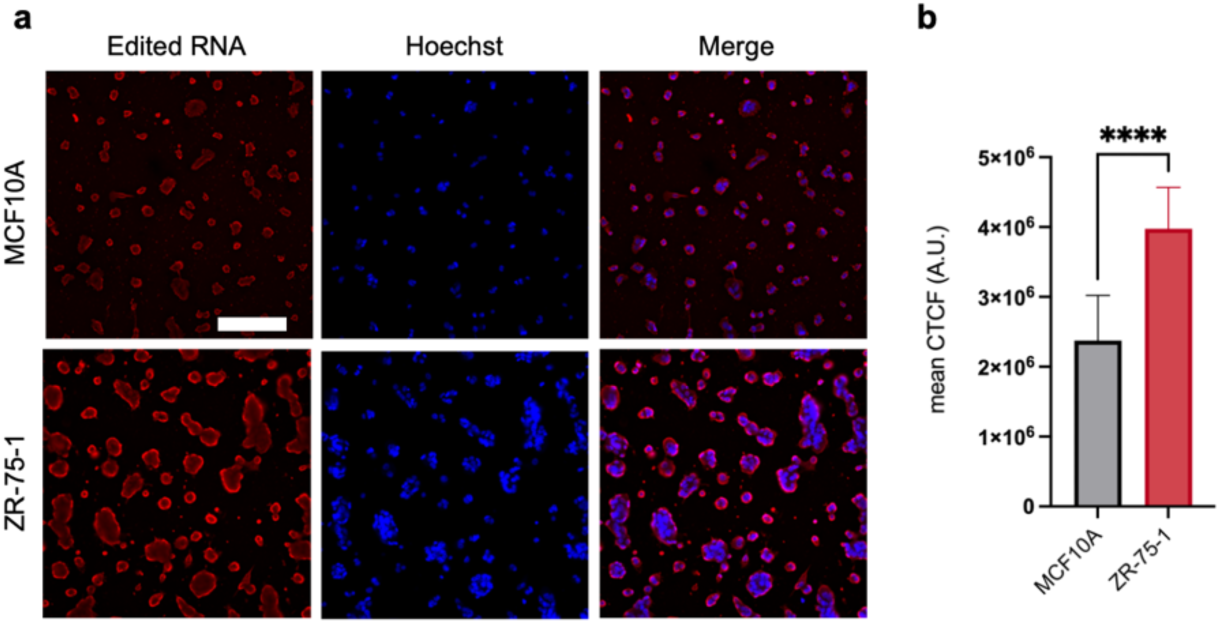
| Identifying hyper-editing in cancerous breast cells. **a**, MCF10A cells (non-malignant) and ZR-75-1 cells (malignant) were fixed and stained for edited RNA (red) and cell nuclei (blue) using EndoVIA. **b,** Quantification of mean corrected total cellular fluorescence (CTCF) in **a**. Data in **a** and **b** are representative of three independent experiments; *n=9* wells from a 96-well plate. Scale bar, 200 μm. Data are shown as mean ± s.d. in arbitrary units (A.U.). Statistical significance was determined by unpaired *t*-test; \*\*\*\**P* < 0.0001.

Having established the ability of EndoVIA to detect cancer-related hyper-editing, we were also curious to explore whether we could detect cancer-related hypo-editing. Kidney cancers such as kidney chromophobe (KICH) and kidney renal papillary cell carcinoma (KIRP) in particular have been identified to display hypo-editing.^60^ Therefore, we selected the G-402 kidney cell line, characterized with renal leiomyoblastoma and an AEI value of 0.8. We also stained HEK293T cells that have an AEI value of 1.1 in parallel as a non-malignant counterpart (Fig. 6a). As anticipated, we were able to detect a decrease in fluorescence in the G-402 cell line (Fig. 6b). We also confirmed that there were comparable amounts of mRNA across each pair of healthy and diseased cell lines, strongly suggesting that differences in fluorescence can be attributed to changes in A-to-I editing (Supplementary Fig. 11).

**Fig. 6.**
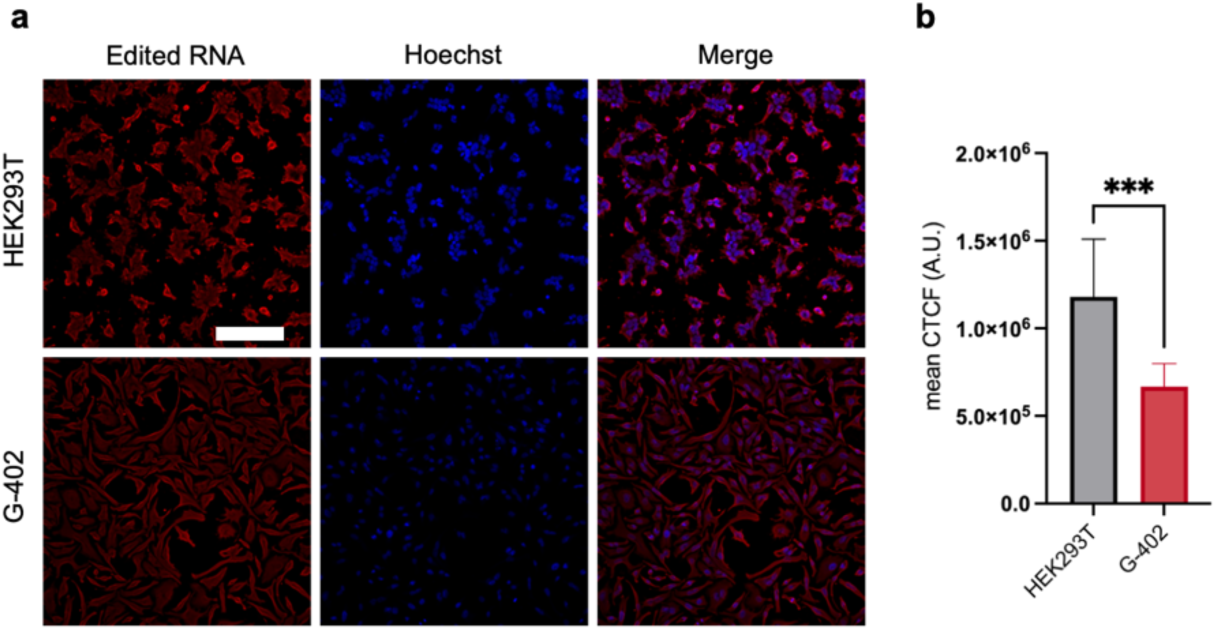
| Identifying hypo-editing in cancerous kidney cells. **a**, HEK293T cells (non-malignant) and G-402 cells (malignant) were fixed and stained for edited RNA (red) and cell nuclei (blue) using EndoVIA. **b,** Quantification of mean corrected total cellular fluorescence (CTCF) in **a**. Data in **a** and **b** are representative of three independent experiments; *n=9* wells from a 96-well plate. Scale bar, 200 μm. Data are shown as mean ± s.d. in arbitrary units (A.U.). Statistical significance was determined by unpaired *t*-test; \*\*\**P* < 0.001.

Having demonstrated that EndoVIA can detect global changes in inosine levels between healthy and cancerous cells, we sought to also capture nuanced shifts in cell-to-cell variation, as cellular heterogeneity is a hallmark of cancer. Using the CTCF values of individual cells across all four cell lines, we employed kernel density estimation to unveil the distribution of fluorescence within each cell population.^61^ There was a notable difference in the distribution patterns that emerged between healthy and diseased cells. Particularly striking was the comparison between HEK2393T and G-402 cells, where we observed a distinct migration of the density peak. In HEK293T cells, a higher density of cells exhibited elevated levels of A-to-I editing, while in G-402 cells the density shifted toward lower CTCF values, signaling an increase in cells displaying reduced levels of editing (Supplementary Fig. 12a). A similar trend was also observed in MCF10A and ZR-75-1 cells, where higher A-to-I editing levels in ZR-75-1 cells displayed a shift in cell density towards higher CTCF values (Supplementary Fig. 12b). These findings aligned with our expectations, reinforcing that EndoVIA is capable of not only detecting widespread changes in A-to-I editing but also uncovering nuances in cellular heterogeneity.

EndoVIA serves as the first method capable of detecting inosine-containing transcripts *in situ*, thus we were curious about its performance against traditional approaches for characterizing A-to-I editing in cells, namely dsRNA antibodies. Given that dsRNA is the primary target of ADAR, the J2 and K1 antibodies have been routinely employed for detecting and enriching ADAR substrates. To put these antibodies to the test, we chose to stain dsRNA with J2 and K1 in HEK293T and G-402 cells and compare to EndoVIA that was completed in parallel (Supplementary Fig. 13). The localization patterns for all three treatments varied across both HEK293T and G-402 cells. Notably, the dsRNA antibodies exhibited a surprising absence of fluorescence in the nucleus, despite this being ADAR’s primary residence for RNA editing. K1, specifically in HEK293T cells, showed localization patterns most comparable to EndoV, demonstrating increased fluorescence near the cell membrane. These observable differences across treatments underscore the unknown binding preferences of J2 and K1 in the context of A-to-I editing status. In comparison, EndoVIA offers a higher degree of confidence in staining edited RNA as it directly detects inosine, providing a higher level of detail and reliable staining approach.

### Detecting Subcellular Localization of Inosine-Containing Transcripts

In addition to the ability to rapidly detect global inosine levels in cell samples, we envisioned that EndoVIA might also enable imaging of the subcellular localization of edited RNAs. The majority of A-to-I editing occurs within the 3’ UTRs of mRNA which heavily influences RNA localization. The loss of an edited 3’ UTR has been shown to completely alter the destination of a given RNA.^26–29^ Despite this evident link between A-to-I editing and RNA localization, many unanswered questions remain due to the lack of available methods. As an example of this, the images of HEK293T cells presented in the figures above show observable disparities in fluorescence signal, with signal seeming to be concentrated near the cellular membrane. In order to more rigorously study these localization patterns and achieve greater resolution, we hypothesized that TIRF illumination coupled with super-resolution microscopy would yield high-quality, nanometric spatial insight from EndoVIA. Thus, we performed the immunostaining on both HEK293T and G-402 cell lines using the EndoVIA protocol.

Imaging of HEK293T cells using dSTORM in TIRF illumination enabled us to observe single EndoV binding events (presumably from individual editing sites) at the membrane (Fig. 7a) and adhesion site (Fig. 7b). We observed edited RNA concentrated at the cellular membrane in contrast to the even distribution throughout the adhesion sight. Interestingly, previous reports have determined that cellular adhesion and motility is partly regulated through RNA localization and translation at focal adhesions.^62,63^ Together, these findings offer new potential insights into the downstream functions of editing, as well as mark the first demonstration that provides spatial distribution of A-to-I editing across the cell at the single-molecule level.

**Fig. 7.**
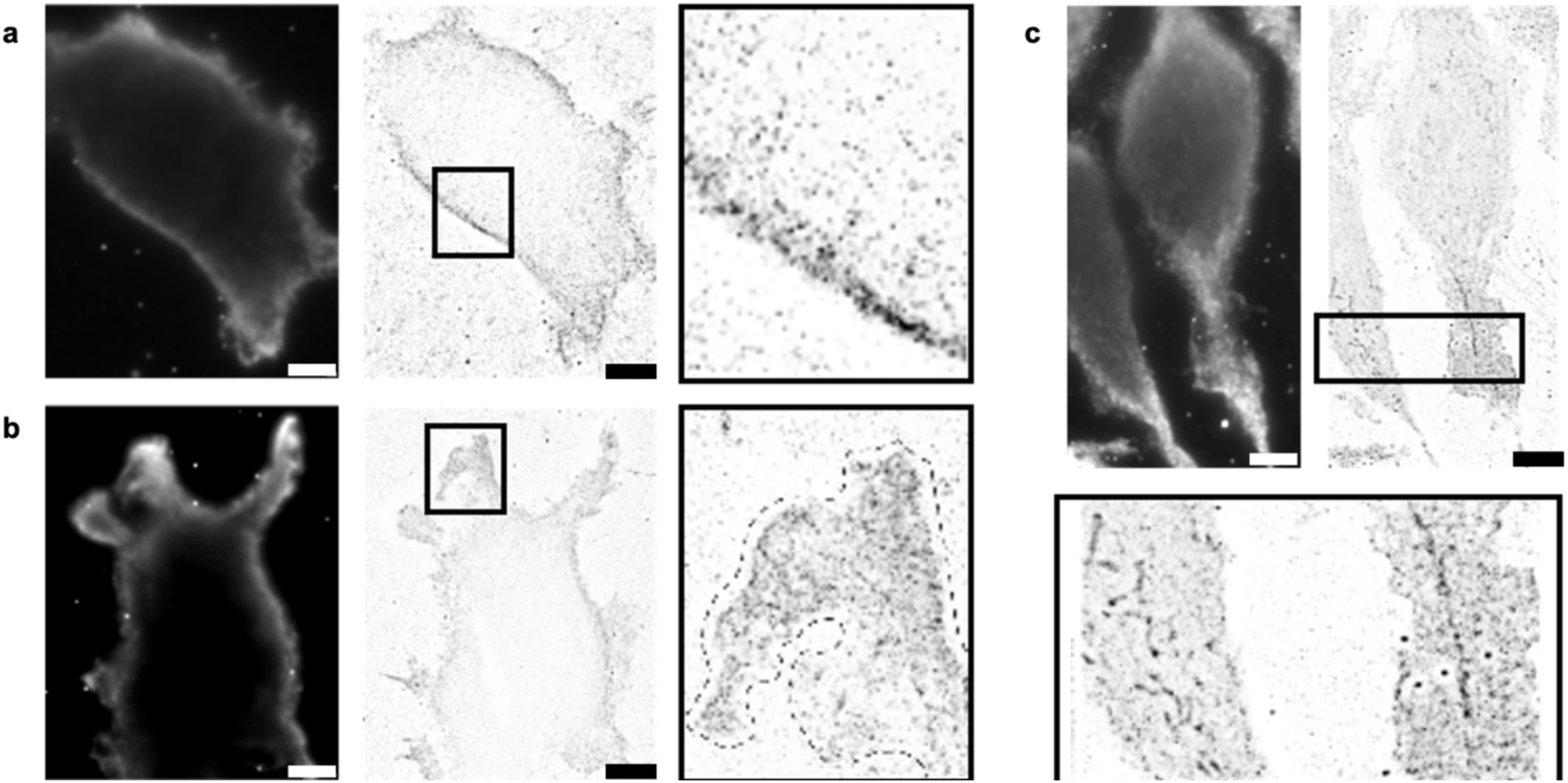
| Super-resolution microscopy of edited RNA using EndoVIA. TIRF (dark images) and TIRF-dSTORM images (light images, whole cell and close-up view of boxed regions respectively) from a, HEK293T cells and b, adhesion site of HEK293T cells and c, G-402 cells labeled with the described immunostaining workflow. Scale bar, 10 µm.

dSTORM TIRF imaging of cancerous G-402 cells show similar localization of edited RNAs at the membrane as well, though less pronounced than in the HEK293T cells. G-402 cells also appear to display more even distribution of edited transcripts throughout the cytoplasm. Interestingly, upon closer inspection, super-resolution imaging of the G-402 cells reveals some clustering in the cytoplasm that may represent subcellular structures for organizing edited RNAs that are not present in HEK293T cells and not visible using standard TIRF or confocal imaging (Fig. 7c). Despite these differences in the localization of edited RNA, ADAR1 localization in both HEK293T cells and G-402 cells remains in the nucleus, hinting at potential underlying mechanisms dictating the localization of edited transcripts that have yet to be discovered (Supplementary Fig. 14). These results not only underscore the power of EndoVIA to lead to new biological discoveries and knowledge, but also establish its compatibility with advanced imaging modalities. These results represent the first example of being able to visualize edited RNAs with nanoscale resolution in the cellular environment. This in turn opens up a wide range of potential explorations that will reveal new links between A-to-I editing and RNA localization and in turn provide novel insights into the role of this important process in development and disease.

## Discussion

A-to-I editing is the most widespread post-transcriptional modification and is essential for multiple biological processes. One way that editing impacts cellular pathways is thought to be through directing the localization of RNAs to subcellular compartments by editing within the 3’ UTR of mRNAs. Although there are a few demonstrations that nuclear retention of specific mRNAs is a result of A-to-I editing, the broader impact of editing on RNA localization is underexplored due to a lack of methods for imaging inosine-containing RNAs in the cellular context. In parallel, dysregulation of A-to-I editing is intricately linked to neurological disorders, autoimmune diseases, and multiple cancers, highlighting its critical role in disease pathogenesis. However, the tremendous potential of editing to serve as a biomarker or therapeutic target is limited by the lack of methods for observing editing directly *in situ*. Specifically, the current gold standard technique of high-throughput RNA sequencing requires RNA extraction and pooling of RNA from multiple cells in a sample. This essentially erases information relating to cell-to-cell variation of global inosine levels and precludes studying the specific subcellular localization of edited RNAs. Moreover, RNA-seq remains limited in throughput, making it poorly suited for high-throughput drug screening campaigns.

Herein, we introduce EndoVIA as the first approach for imaging and quantifying the full breadth of inosine-containing transcripts in cells. Key to the development of this technique is our repurposing of EndoV to act as an “anti-inosine antibody,” which in turn enables us to develop a protocol analogous to immunofluorescence staining that is aimed at detecting edited RNAs *in situ*. We have validated our approach using cells having varying editing levels and demonstrate the ability to detect biologically relevant differences in inosine between healthy and cancerous cell lines. We also show that EndoVIA is compatible with super-resolution microscopy techniques to image edited RNAs at nanoscale resolution and provide previously unattainable insight into the subcellular localization and organization of edited transcripts.

We envision that the availability of EndoVIA will open up numerous avenues for studying RNA editing and harnessing this important process for diagnostics and therapeutics. For example, efforts are underway in our laboratory to further elaborate and refine EndoVIA to enable high-throughput phenotypic screening for the discovery of novel small molecules to regulate A-to-I editing. Additionally, the ability to image the subcellular localization of edited RNAs can be directed toward probing a wide range of biological questions regarding how this localization is impacted by cell type, disease state, and external stimuli. Thus, this first-in-class approach for imaging A-to-I editing at the cellular level is expected to not only advance research in our own laboratory but also empower other groups studying A-to-I editing and in turn advance both the basic science and translational potential of this important biological process.

## Acknowledgements

This work was supported by the National Institutes of Health (R35GM144075 to J.M.H.). The content is solely the responsibility of the authors and does not necessarily reflect the official views of the National Institute of Health. This research project was supported in part by the Emory University Integrated Cellular Imaging Microscopy Core and the Washington University in St. Louis Genomic Technology Access Center. We would like to thank Dr. Charles Rice for gifting us the ADAR1 KO cell line. We would also like to thank Kevin Kaifer for guidance with image acquisition.

## Methods

### Materials

Methanol, 4% paraformaldehyde, ethanol, glyoxal, acetic acid, Triton X-100, Tween 20, tris hydrochloride, calcium chloride, sodium chloride, cholera toxin, TE buffer, and EDTA were purchased from Sigma Aldrich. Nuclease-free water, Hoechst 33342, and 6-well plates were purchased from Thermo Fisher Scientific.

### Cell Culture and Transfection

The G-402 cell line (ATCC) was cultured in McCoy’s 5a Medium Modified (ATCC) supplemented with 10% fetal bovine serum (Gibco) and 1% penicillin-streptomycin (Gibco). The HEK293T cell line (ATCC) and the HEK293T ADAR1 KO cell line (gifted by Dr. Charles Rice) was cultured in Dulbecco’s Modified Eagle’s Medium (Gibco) supplemented with 10% fetal bovine serum and 1% penicillin-streptomycin. The MCF10A cell line (ATCC) was cultured in MEBM Basal Medium (Lonza) supplemented with the included additives, excluding GA-1000, and 100 ng mL^-1^ cholera toxin. The ZR-75-1 cell line (ATCC) was cultured in RPMI-1640 Medium (ATCC) supplemented with 10% fetal bovine serum and 1% penicillin-streptomycin. All cell lines were cultured at 37°C in a humidified incubator with 5% CO_2_. For FISH and immunofluorescence experiments, cells were seeded and incubated for 48 hours at 37°C in a humidified incubator with 5% CO_2_. For transfection, cells were seeded, and 24 hours post seeding (∼70% confluent), cells were transfected with increasing amounts of ADAR1-GFP plasmid (Addgene) or coilin-GFP plasmid (Addgene) using Opti-MEM Reduced Serum Medium (Gibco) and lipofectamine 3000 (Invitrogen) following the manufacturer’s protocol. Cells were then incubated for 48 hours at 37°C in a humidified incubator with 5% CO_2_ before completing subsequent immunostaining.

### tRNA and GAPDH Fluorescence *In Situ* Hybridization (FISH)

For tRNA FISH, cells were seeded as previously described. Wells were then fixed with 100% methanol or 4% paraformaldehyde. Fixative reagents were then removed, and wells were washed with 1X PBS (Invitrogen). Cells were permeabilized in 0.1% Triton X-100 in 1X PBS for 10 minutes. The Stellaris RNA FISH hybridization buffer, Wash Buffer A, and Wash Buffer B were prepared according to the manufacturer’s instructions (Stellaris). The 1X PBS was then replaced with the prepared Stellaris RNA FISH Wash Buffer A and incubated for 5 minutes. tRNA FISH probes (IDT, Supplementary Table 1) were dissolved in TE buffer (10 mM tris hydrochloride, 1 mM EDTA, pH 8.0) at a concentration of 100 μM and then diluted in the prepared hybridization buffer to a concentration of 250 nM. Wash Buffer A was then replaced with the diluted probes, the plate was sealed to prevent evaporation, and incubated in the dark at 37°C overnight. The probes were removed and replaced with Wash Buffer A and incubated in the dark at 37°C for 30 minutes. Wash Buffer A was removed and replaced with Wash Buffer B and incubated for 5 minutes. Wash Buffer B was exchanged for 1X PBS and cells were imaged. For GAPDH FISH, cells were seeded as previously described. Wells were then fixed with 100% methanol. The methanol was removed, and cells were washed with 1X PBS. Wells were treated with either 3%, 6%, 12%, or 30% glyoxal solution and incubated at 50°C for 1 hour. The glyoxal solutions were removed and wells were washed twice with 1X PBS. Cells were then permeabilized in 0.1% Triton X-100 in 1X PBS for 10 minutes. GAPDH FISH was completed by using Human GAPDH FISH Probes (Stellaris) and the previously described protocol for tRNA FISH. Cells were then imaged.

### Immunofluorescence

Nup153 and β-actin were immunstained in the following manner. Cells were cultured and seeded as previously described. Cells were then fixed in 100% methanol. The methanol was removed, and wells were washed twice with 1X PBS. A 4% glyoxal denaturing solution was prepared and incubated at 37°C for 1 hour. Wells were then washed twice with 1X PBS. The 1X PBS was replaced with a 0.1% Triton X-100 in 1X PBS permeabilization solution and incubated for 10 minutes followed by two washes in 1X PBS. Cells were then incubated in a blocking solution containing 3% bovine serum albumin (Gibco), 0.1% Triton X-100, and 0.1% Tween 20 in 1X PBS for 1 hour. Wells were washed twice in 1X PBS and then incubated in anti-Nup153 antibody (Abcam) or anti-β-actin antibody (Invitrogen) in blocking buffer for 1 hour. The primary antibody solutions were removed, and wells were washed three times in 1X PBS. Finally, a staining solution comprised of Hoechst nuclear dye and goat anti-rabbit Alexa Fluor 647 (Invitrogen, Nup153) or goat anti-rabbit Alexa Fluor 488 (Invitrogen, β-actin) diluted in blocking buffer were added and incubated for 1 hour in the dark. Wells were then washed in 1X PBS three times and imaged. To assess the cellular morphology under varying glyoxal conditions, β-actin was stained in the following manner. Cells were cultured and seeded as previously described. Cells were then immunostained as previously described, with the exception that multiple glyoxal concentrations were tested (4%, 12%, 30%). ADAR1 and dsRNA were stained in the following manner. Cells were cultured and seeded as previously described. Cells were then fixed in 4% paraformaldehyde and then washed, permeabilized, and blocked as previously described for Nup153 and β-actin. After washing with 1X PBS, cells were incubated in primary antibody solutions consisting of anti-dsRNA J2 antibody (Sigma Aldrich), anti-dsRNA K1 antibody (Nordic MUbio), or anti-ADAR1 antibody (Atlas Antibodies) in blocking buffer and incubated for 1 hour. The primary antibody solutions were removed, and wells were washed three times in 1X PBS. Finally, a staining solution comprised of Hoechst nuclear dye and goat anti-mouse Alexa Fluor 647 (Invitrogen, J2 and K1) or goat anti-rabbit Alexa Fluor 647 (Invitrogen, ADAR1) diluted in blocking buffer were added and incubated for 1 hour in the dark. Wells were then washed in 1X PBS three times and imaged.

### EndoVIA

Cells were seeded as previously described. Cells were then fixed in methanol and subsequently washed twice with 1X PBS. The 1X PBS was removed and the optimized glyoxal denaturing solution was placed on cells and incubated for 1 hour. Cells were then incubated with a 0.1% Triton X-100 in 1X PBS permeabilization solution for 10 minutes followed by two washes in 1X PBS. Cells were incubated in a blocking solution containing 3% bovine serum albumin, 0.1% Triton X-100, and 0.1% Tween 20 in 1X PBS for 1 hour. Wells were then washed twice with 1X PBS. Cells were incubated with 1:50 Endonuclease V (New England BioLabs) in a calcium containing blocking buffer, or the calcium containing blocking buffer alone as a negative control, for 1 hour. Cells were then washed three times in a calcium containing wash buffer, incubating for 5 minutes each time. Cells were then incubated for 1 hour with anti-MBP antibody (New England BioLabs) solution diluted in calcium containing blocking buffer. Cells were washed three times, with 5-minute incubation periods. Finally, a staining solution containing goat anti-mouse Alexa Flour 647 and Hoechst nuclear dye in calcium containing blocking buffer was incubated in the wells for 1 hour in the dark. Cells were washed three times for 5 minutes each time. Cells were then imaged. The EndoV concentration used for the EndoVIA workflow was determined by following the previously described protocol using a range of EndoV and MBP (Novus Biologicals) concentrations (1:4000-1:25). The calcium dependence of EndoVIA was confirmed by treating wells with 5 mM EDTA and incubating for 1 hour following EndoVIA and then imaged.

### mRNA Quantification

Cells were seeded in 6-well plates and incubated at 37°C in a humidified incubator with 5% CO_2_ for 24 hours before harvesting for RNA. Cells were lysed and mRNA was isolated and purified using the Magnetic mRNA Isolation Kit (New England BioLabs). Resulting purified mRNA was quantified using Nanodrop.

### Microscopy and Image Analysis

Stained cells were imaged using a Nikon Spinning Disk for widefield microscopy with a 20x air objective and confocal microscopy with a 60x oil objective. Laser excitation at 405 nm was used to image Hoechst 33342; excitation at 640 nm was used to image Alexa Fluor 647; excitation at 488 nm was used to image ADAR-GFP; excitation at 560 nm was used to image Quasar570 labeled GAPDH FISH probes. Gain and exposure settings for each laser were optimized to achieve sufficient fluorescence and minimize oversaturation. The resulting images were analyzed to determine the fluorescence of each cell using FIJI. The area and integrated density were measured for each cell and then the Corrected Total Cellular Fluorescence was calculated as follows: 𝐶𝑇𝐶𝐹 = 𝑖𝑛𝑡𝑒𝑔𝑟𝑎𝑡𝑒𝑑 𝑑𝑒𝑛𝑠𝑖𝑡𝑦 − (𝑐𝑒𝑙𝑙 𝑎𝑟𝑒𝑎 × 𝑚𝑒𝑎𝑛 𝑏𝑎𝑐𝑘𝑔𝑟𝑜𝑢𝑛𝑑 𝑓𝑙𝑢𝑜𝑟𝑒𝑠𝑐𝑒𝑛𝑐𝑒).

### Alu Editing Index (AEI)

Cells were seeded in 6-well plates and incubated at 37°C in a humidified incubator with 5% CO_2_ for 48 hours. Cells were lysed and total RNA was isolated and purified using the Monarch Total RNA Miniprep Kit (New England BioLabs). This purified RNA was then used to prepare sequencing libraries with the Tru-Seq Stranded with RiboZero Gold (Human/Mouse/Rat) Kit. Standard 8-bp i5 and i7 Illumina index barcode and adapters were added to each library. Libraries were sequence using a NovaSeq X Plus 300 cycles (Illumina) to produce paired end 150-bp reads (approximately 30M reads per sample). Raw FASTQ files were trimmed using Trimmomatic1 with the parameter HEADCROP:3 to remove the first 3 bp.^64^ FASTQC (https://www.bioinformatics.babraham.ac.uk/projects/fastqc/) was then used to check read quality (PHRED33) after data trimming. Reads were next aligned to the human reference genome hg38 using STAR 2.5.23 with the additional parameter --outFilterMatchNminOverLread 0.95 to detect A-to-I editing.^65^ The resulting .bam files were sorted, and duplicates were removed using Samtools 1.35.^66^ Finally, the RNA editing indexer package was used with the default settings to calculate the AEI for each sample.^12^

### Kernel Density Estimation

This analysis utilized Python with the pandas, seaborn, and matplotlib libraries to create Kernel Density Estimation (KDE) plots from the individual CTCF values of each cell. The CTCF values underwent preprocessing and were loaded into a pandas DataFrame. KDE plots were then generated using the seaborn library’s kdeplot () function. The python code was written and executed using Visual Studio Code.

### dSTORM super-resolution Setup and Imaging

dSTORM images were acquired on a home-built setup based on a Nikon Ti2 Eclipse microscope body operated with Micro-Manager 2.0. Excitation was performed by a 180mW 638nm laser (Cobolt 06-MLD, Coherent) to which the beam was circularly polarized (ThorLabs), expanded (GBE10-A, ThorLabs) and focused at the back focal plane of the objective (SR HP TIRF, 100X, 1.49NA, Nikon) by an achromatic lens (f=300mm, AC254-300-A, ThorLabs) mounted on a motorized translating stage (KMTS25E, ThorLabs) to achieve TIRF illumination. A filter cube (TRF89901v2, Chroma) containing excitation and emission filters ensure spectral filtering of both light paths. Emission was detected by a sCMOS camera (Orca Fusion, Hamamatsu) with 2×2 binning, resulting in effective pixel size of 130 nm. Prior to imaging, cells were treated with dSTORM imaging buffer (GLOX) consisting of 0.56 mg.mL-1 Glucose Oxidase (Sigma), 34 µg.mL-1 Catalase (Sigma), 50 mM Tris (pH 8.00), 10 mM NaCl, 10% glucose (w/v) and 143 mM β-mercaptothanol. For imaging, a stack of 20,000 images were taken per field at maximum laser power. The first 1,000 images were discarded and the remaining stacks were analyzed by ThunderSTORM imageJ plugin. Microscope’s Perfect Focus System (PFS, Nikon) compensate for Z drift, while 100nm fiducial marker (TetraSpeck, Invitrogen) were added post-fixation to compensate for XY drift.

### Statistics

A minimum of three biological replicates were completed for each experiment. Statistical analyses were completed using GraphPad Prism 10 and all values and error bars indicate the mean ± s.d. unless otherwise noted. For comparison of two independent groups, an unpaired *t*-test was performed. For multiple comparisons, one-way ANOVA was performed.

