## Supplemental Figures for "Spatial visualization of A-to-I Editing in cells using Endonuclease V Immunostaining Assay (EndoVIA)"

### Supplemental Information

#### Table of Contents

**S Table 1.** tRNA FISH Probes S2

**S Fig. 1.** Fluorescence *in situ* hybridization (FISH) of tRNA using different fixatives S3

**S Fig. 2.** Detection of Nup153 and β-actin using modified immunofluorescence workflow S4

**S Fig. 3.** Non-specific binding of MBP of the EndoV-MBP fusion protein S5

**S Fig. 4.** Optimizing glyoxal concentration using *GAPDH* fluorescence *in situ* hybridization (FISH) S6

**S Fig. 5.** Cellular morphology of cells treated with varying glyoxal concentrations using β-actin S7

**S Fig. 6.** Optimizing EndoV concentration S8

**S Fig. 7.** EndoV stained cells treated with EDTA S9

**S Fig. 8.** Immunostaining ADAR1 S10

**S Fig. 9.** AEI Values of WT and ADAR1 KO HEK293T cells S11

**S Fig. 10.** Quantifying A-to-I editing in HEK293T cells transfected with coilin-GFP S12

**S Fig. 11.** Quantifying mRNA in non-malignant and malignant cell lines S13

**S Fig. 12.** Cellular heterogeneity in non-malignant and malignant cell lines S14

**S Fig. 13.** Immunostaining dsRNA and edited RNA in healthy and diseased Cells S15

**S Fig. 14.** Immunostaining ADAR1 in HEK293T and G-402 cells S16

| tRNA Ser AGA | |
|---|---|
| **Probe 1** | 5’ – AA+C CA+C TC+G GC+C AC+G AC+T AC/3AlexF488N/ – 3’ |
| **Probe 2** | 5’ – CGC +GGG +GAA +ACC +CCA +ATG +GAT +TTC /3AlexF88N/ –3’ |

**Supplementary Table 1 | tRNA Fluorescence *in situ* hybridization (FISH) probes.** The following locked nucleic acid (LNA) FISH probes were used to detect tRNA Ser AGA. LNA bases are denoted with (+) following the nucleobase and probes are labeled with Alexa Fluor 488 (3AlexF88N).

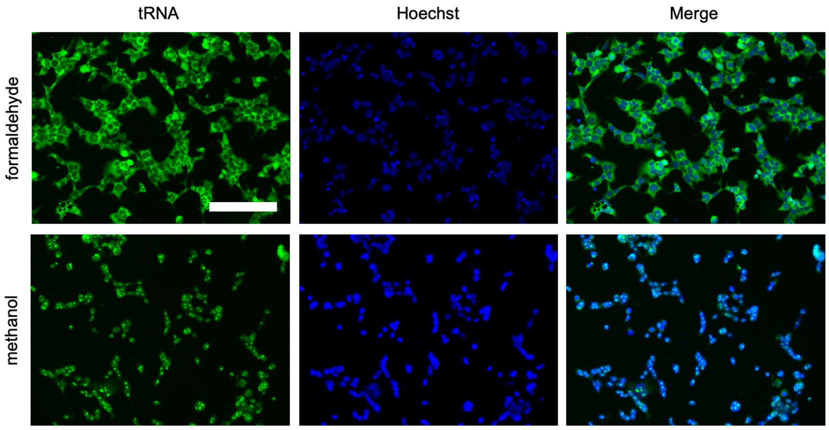


**Supplementary Fig. 1 | Fluorescence *in situ* hybridization (FISH) of tRNA using different fixatives.** HEK293T cells were fixed with either formaldehyde or methanol and stained for tRNA (green) and cell nuclei (blue). Data are representative of three independent experiments; *n=3* wells from a 96-well plate. Scale bar, 200 μm.

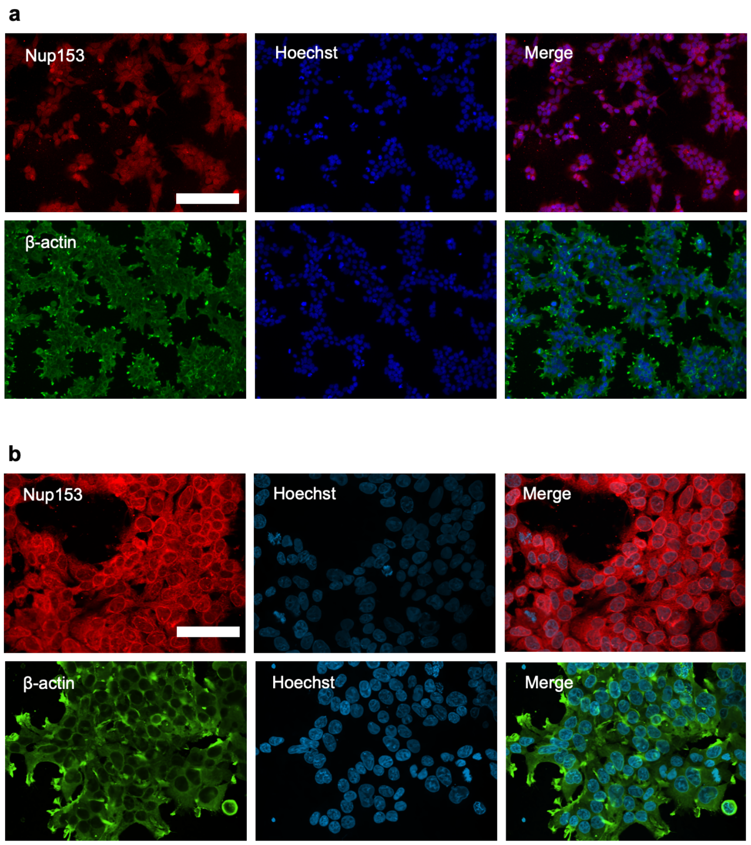


**Supplementary Fig. 2 | Detection of Nup153 and β-actin using modified immunofluorescence workflow. a,** HEK293T cells were fixed and stained for Nup153 (red) or β-actin (green) and cell nuclei (blue). Images were taken using widefield microscopy. **b,** HEK293T cells were prepared as previously described in **a** and imaged using confocal microscopy. Data are representative of three independent experiments; *n=3* wells from a 96-well plate. Scale bar, **a** 200 μm and **b** 50 μm.

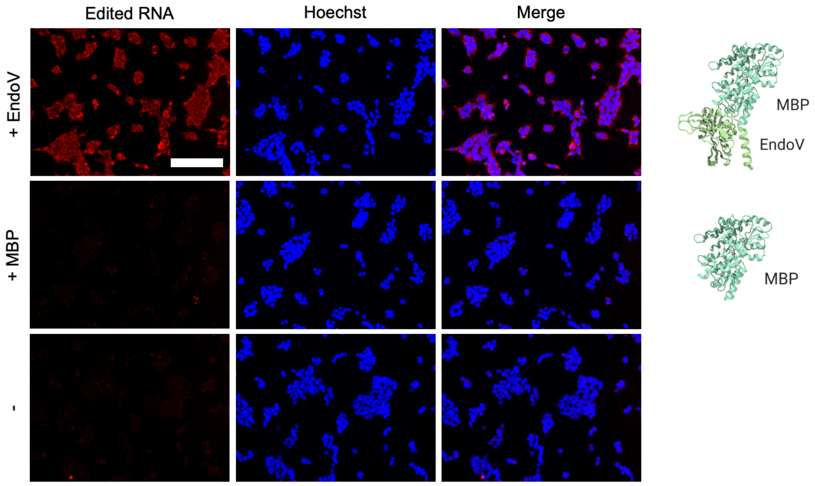


**Supplementary Fig. 3 | Non-specific binding of MBP of the EndoV-MBP fusion protein.** HEK293T cells were fixed and stained for edited RNA (red) and cell nuclei (blue) using EndoV-MBP fusion protein (+ EndoV), MBP alone (+ MBP), or neither (-). Data are representative of three independent experiments; *n*=3 wells from a 96-well plate. Scale bar, 200 µm.

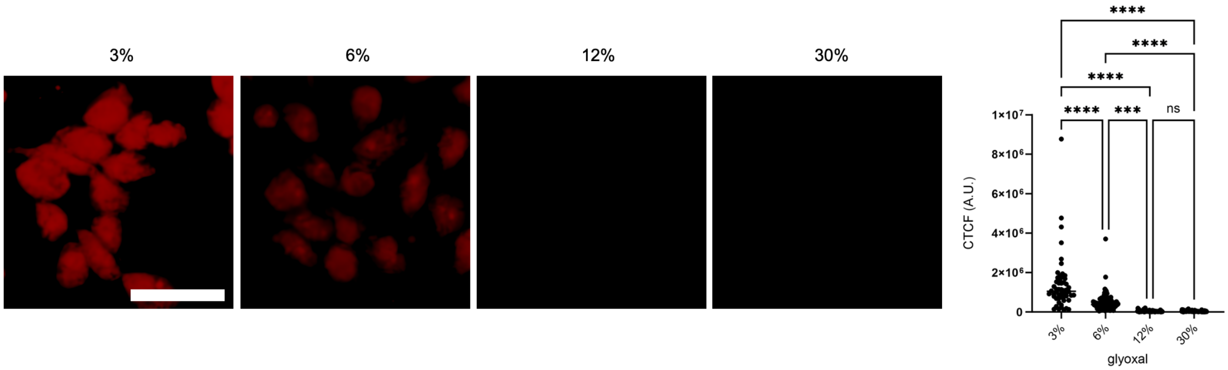


**Supplementary Fig. 4 | Optimizing glyoxal concentration using *GAPDH* fluorescence *in situ* hybridization (FISH).** HEK293T cells were fixed and treated with increasing amounts of glyoxal (3%, 6%, 12%, 30%) and stained for *GAPDH* mRNA using FISH probes (red) and quantified for mean corrected total cellular fluorescence (CTCF). Data are representative of three independent experiments; *n=50* cells. Scale bar, 50 µm. Data are shown as CTCF per cell in arbitrary units (A.U.). Statistical significance was determined by one-way ANOVA; not significant (ns), ***$P$ < 0.001, ****$P$ < 0.0001.

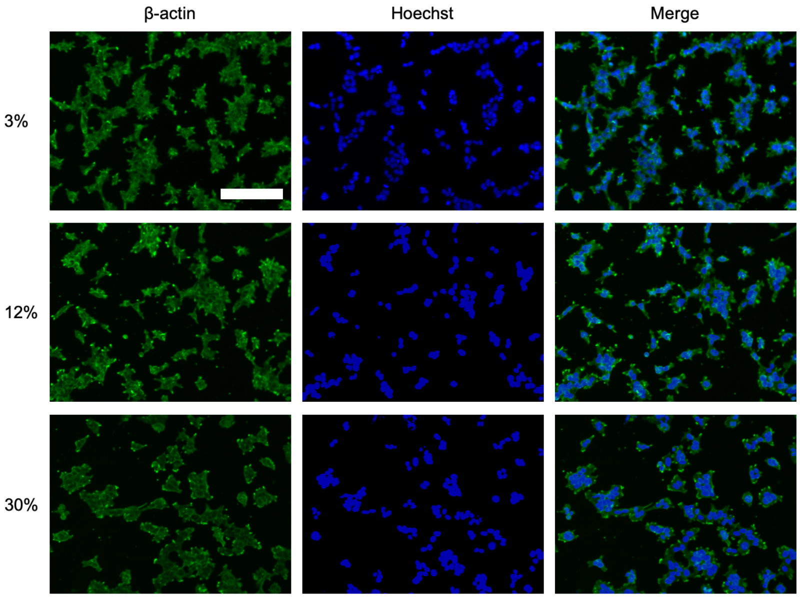


**Supplementary Fig. 5 | Cellular morphology of cells treated with varying glyoxal concentrations using β-actin.** HEK293T cells were fixed and treated with increasing amounts of glyoxal (3%, 12%, 30%) and stained for β-actin (green) and cell nuclei (blue). Data are representative of three independent experiments. Scale bar, 200 μm.

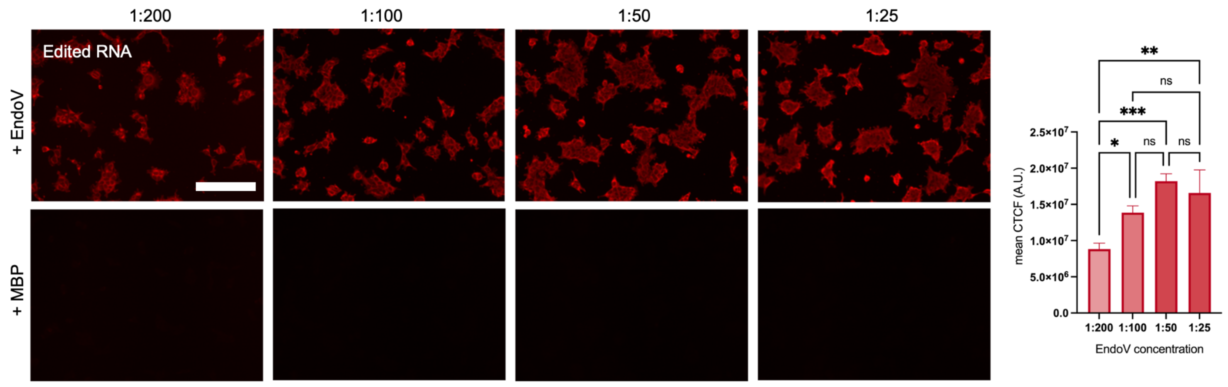


**Supplementary Fig. 6 | Optimizing EndoV concentration.** HEK293T cells were fixed and stained for edited RNA (red) with increasing amounts of EndoV or MBP and quantified for mean corrected total cellular fluorescence (CTCF). Data are representative of three independent experiments; *n=3* wells from a 96-well plate. Scale bar, 200 µm. Data are shown as mean ± s.d. in arbitrary units (A.U.). Statistical significance was determined by one-way ANOVA; not significant (ns), **P* < 0.05, ***P* < 0.01, ****P* < 0.001.

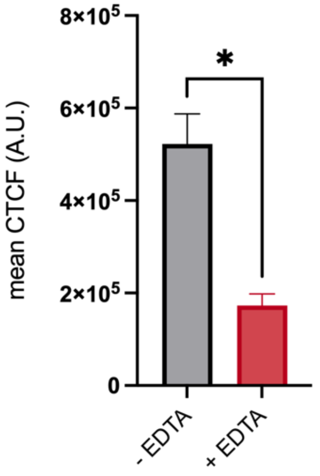


**Supplementary Fig. 7 | EndoV stained cells treated with EDTA.** HEK293T cells were fixed and stained for edited RNA, treated with EDTA, and quantified for mean corrected total cellular fluorescence (CTCF). Data are representative of three independent experiments; *n=3* wells from a 96-well plate Data are shown as mean ± s.d. in arbitrary units (A.U.). Statistical significance was determined by unpaired *t*-test; $^*P < 0.05$.

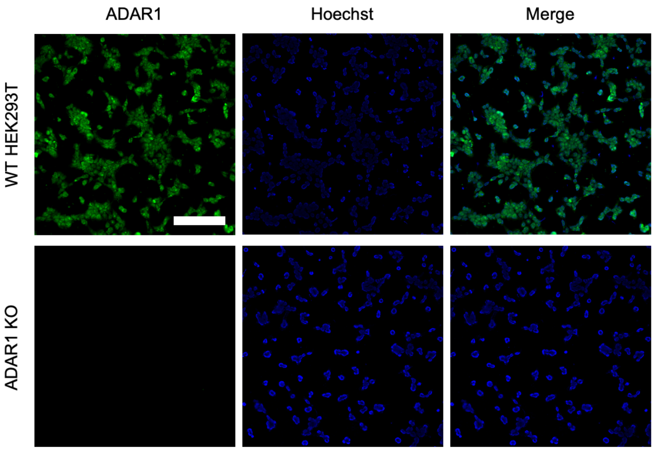


**Supplementary Fig. 8 | Immunostaining ADAR1.** WT HEK293T cells and ADAR1 KO cells were fixed and stained for ADAR1. Data are representative of three independent experiments; *n=3* wells from a 96-well plate. Scale bar, 200 μm.

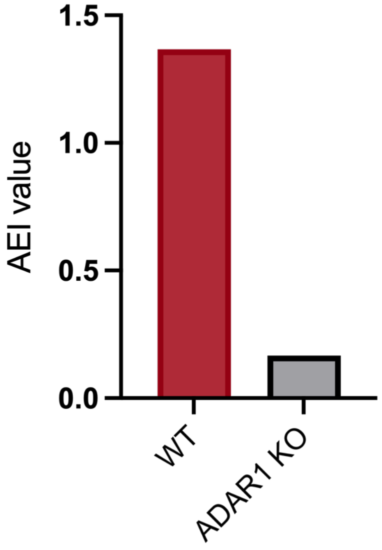


**Supplementary Fig. 9 | AEI Values of WT and ADAR1 KO HEK293T cells.** Total RNA was isolated, purified, and sequenced. Resulting datasets were then trimmed, aligned, and sorted to calculate the AEI values.

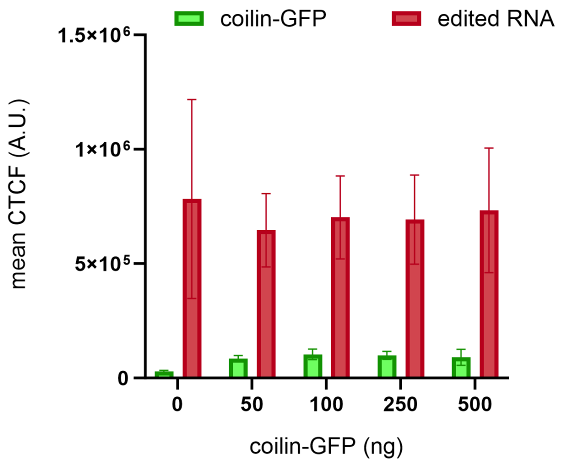


**Supplementary Fig. 10 | Quantifying A-to-I editing in HEK293T cells transfected with coilin-GFP.** HEK293T cells were transfected with increasing amounts of coilin-GFP plasmid (0-500 ng), fixed, and stained for edited RNA (red) using EndoVIA. Coilin-GFP and edited RNA fluorescence were quantified for mean corrected total cellular fluorescence (CTCF). Data are representative of three independent experiments; *n=3* wells from a 96-well plate. Data are shown as mean ± s.d. in arbitrary units (A.U.).

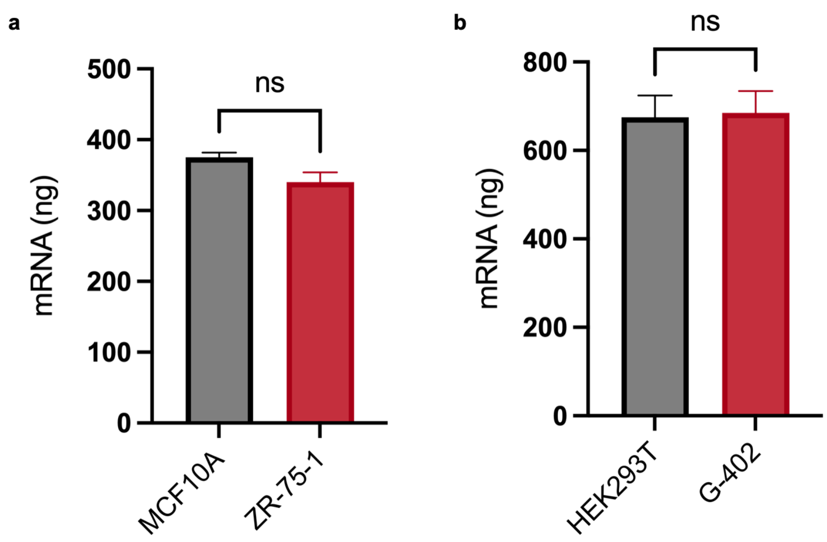


**Supplementary Fig. 11 | Quantifying mRNA in non-malignant and malignant cell lines.** The mRNA from MCF10A, ZR-75-1, HEK293T, and G-402 cell lines were isolated and quantified. Data are representative of three independent experiments; *n=3* wells from a 6-well plate. Data are shown as mean ± s.d. Statistical significance was determined by unpaired *t*-test; not significant (ns).

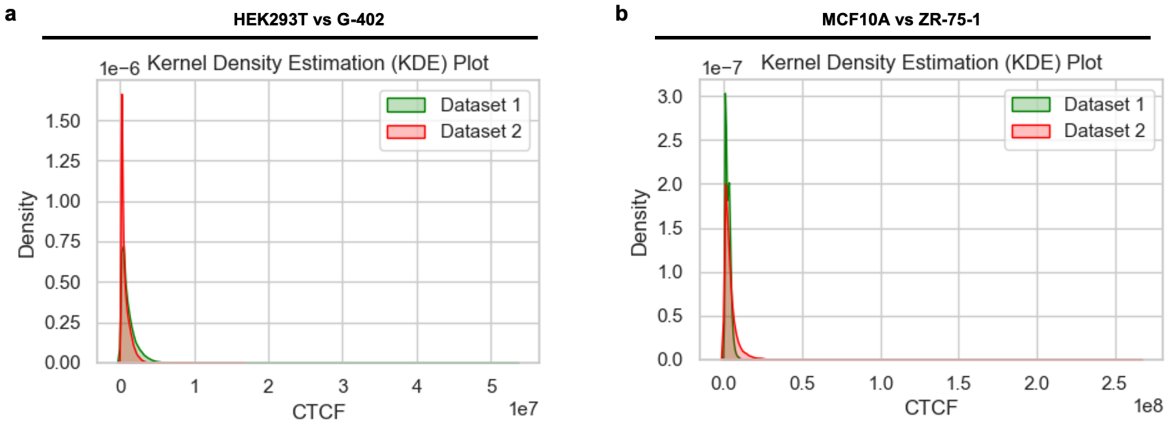


**Supplementary Fig. 12 | Cellular heterogeneity in non-malignant and malignant cell lines.** The kernel density estimation was determined for **a,** HEK293T (green) and G-402 (red) cells and **b,** MCF10A (green) and ZR-75-1 (red) cells using the CTCF values of each individual cell. Data are representative of three independent experiments; *n=9* wells from a 96-well plate.

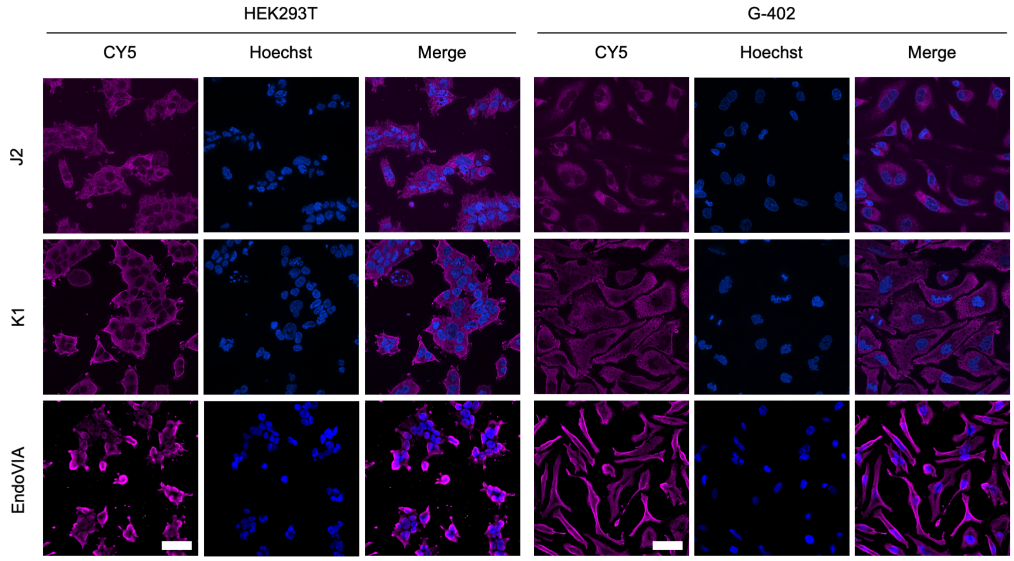


**Supplementary Fig. 13 | Immunostaining dsRNA and edited RNA in healthy and diseased cells.** Fixed HEK293T and G-402 cells were immunostained for dsRNA (J2, K1) or edited RNA (EndoVIA). Data are representative of three independent experiments; *n=3* wells from a 96-well plate. Scale bar, 50 µm.

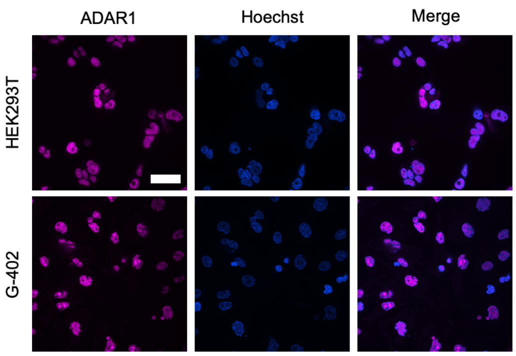


**Supplementary Fig. 14 | Immunostaining ADAR1 in HEK293T and G-402 cells.** Fixed HEK293T and G-402 cells were immunostained for ADAR1. Data are representative of three independent experiments; *n=3* wells from a 96-well plate. Scale bar, 50 µm.
